# Dynamic, non-contact 3D sample rotation for microscopy

**DOI:** 10.1101/177733

**Authors:** Frederic Berndt, Gopi Shah, Jan Brugués, Jan Huisken

## Abstract

*In vivo* imaging of growing and developing samples requires a dynamic adaptation of the sample orientation to continuously achieve optimal performance. Here, we present how, after the injection of magnetic beads, a sample can be freely positioned by applying a magnetic field. We demonstrate its performance for zebrafish on an epi-fluorescence microscope and on a light sheet system for superior multi-view acquisition.

## Main text

Imaging biological samples with light microscopy often requires a certain sample orientation for obtaining ideal image quality. For example, in developing embryos absorbing and scattering tissues such as pigments, eyes or yolk can obscure the area of interest in all but one orientation. Since the sample is embedded typically prior to the experiment, its orientation is fixed. Thereby the imaging is limited to a certain region for the whole duration of the experiment and the orientation cannot be adapted according to the sample’s development during *in vivo* experiments. Hence, to study living organisms with optimal resolution, a technique to dynamically adjust sample orientation in the microscope is needed.

Sample orientation techniques have been developed for high-throughput applications using microfluidic systems ^1^ and for analysis of expression patterns in zebrafish larval brains by manually turning the sample ^2^. However, these methods still lack adaptive reorientation of the sample during the experiment. In some implementations of light-sheet microscopy (or SPIM ^3^) a certain degree of adaptive reorientation of the sample is offered by a vertical rotational axis. However, this uniaxial rotation is insufficient for total control over three-dimensional orientation. Here, we present a non-contact sample orientation technique for living samples. After the injection of magnetic beads, the sample can be freely positioned in 3D during the experiment by applying a magnetic field. We use this technique in freely developing zebrafish embryos and larvae.

Optical methods have been successfully used to position and orient single cells and small worm embryos (*Pomatoceros lamarckii*, 60µm) for light microscopy ^4–7^, but the forces are not sufficient to position a millimeter-sized zebrafish embryo or other model organisms common in developmental biology. To overcome this limitation, we used magnetic forces to orient an early zebrafish embryo within its chorion.

We injected superparamagnetic beads into the yolk using a microinjection needle (**Figure 1a**). Injections were performed between 2.5 hpf and 4 hpf and the injection needle was inserted from either the vegetal pole or the lateral side and beads were deposited close to the yolk membrane. We found that performing the injections at low pressure (10-15 psi) and long injection duration (100-150 ms) avoided the dispersion of beads. By subsequently applying a strong constant magnetic field with a permanent magnet, the beads were attracted and clumped, preventing single beads from moving through the yolk. We used superparamagnetic beads, exhibiting magnetic properties only in the presence of a magnetic field ^8^ since no residual force should be present in the sample after orientation. The beads were chosen to be small enough to be injected without damaging the zebrafish, but large enough that viscous forces prevented them from moving through the yolk. We found that a bead size of 2.8 µm allowed for a rotation of the sample with moderate magnetic fields without translation of the beads within the zebrafish chorion. The applied force on the fish is proportional to the number of injected beads hence a higher volume of beads is favorable. A volume as low as 1 nl bead solution corresponding to about 15 ng of beads or about 1000 beads was sufficient to rotate the embryo without damaging the fish.

**Figure 1.**
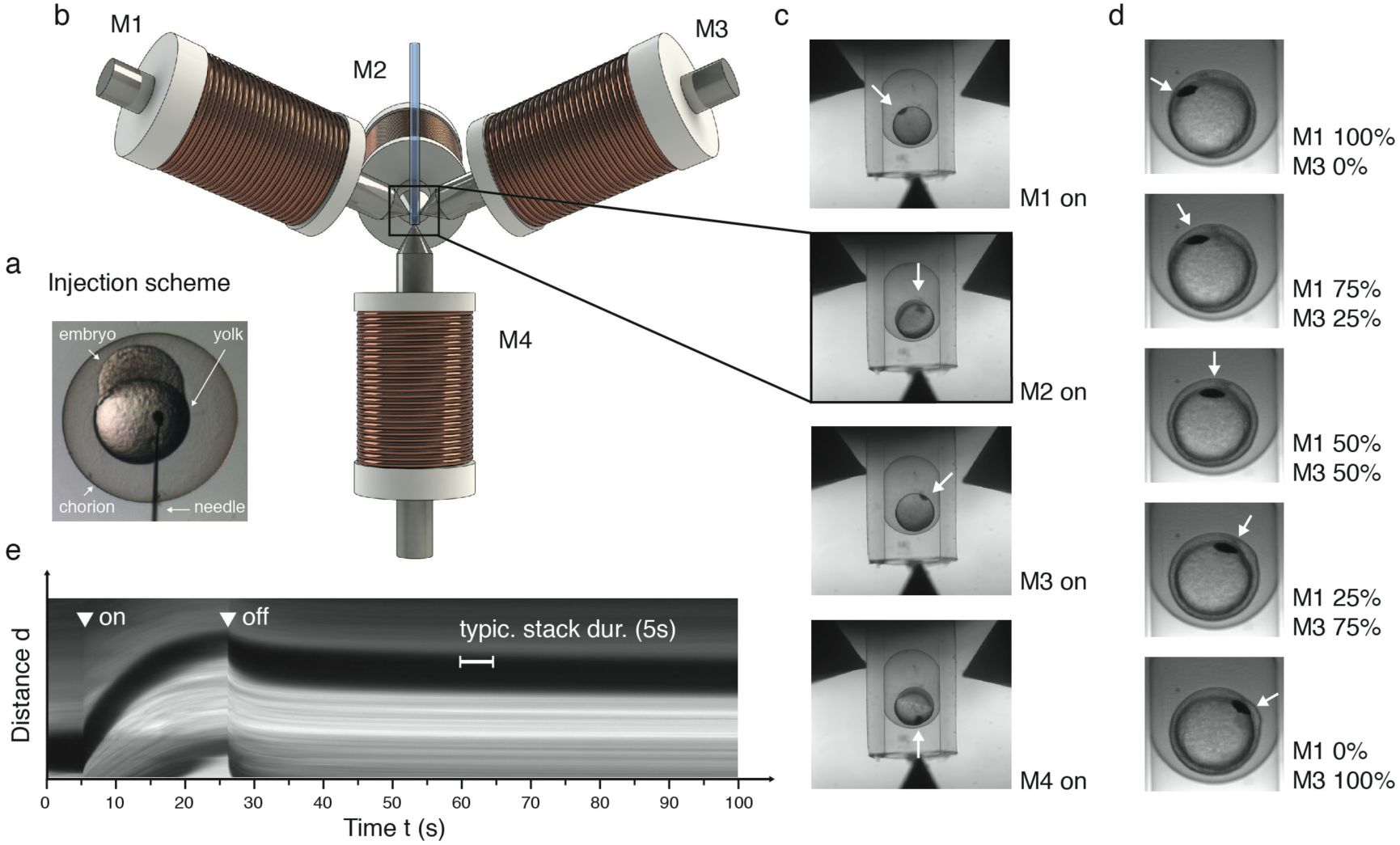
Non-contact positioning technique for embryo imaging from various sides. (***a***) Four electromagnets (M1, M2, M3, M4) in a tetrahedral geometry were assembled around the sample tube. (***b***) Bright-field images of a zebrafish embryo encapsulated in the sample tube. The embryo was oriented by applying a magnetic field by magnet M1, M2, M3 and M4. (***c***) The zebrafish embryo was rotated continuously from one magnet (M1) to the other one (M3) by changing the ratio of the applied currents between two magnets. (***d***) The magnetic beads had been injected at 4hpf into the yolk of the zebrafish embryo. (***e***) Kymograph of a reorienting zebrafish embryo. The arrow heads indicate the time-point when the electromagnet was switched on and off again. The rotation of the zebrafish embryo from one magnet position to the next one (109.5°) took less than 30s. After a retraction of less than 10 s the embryo remained stable in its settle position for over a minute longer than the typical duration of stack (few seconds) in a SPIM setup.

We monitored the injected embryos for 4 days and found no visible delay or defect in development when compared to non-injected wildtype larvae **(Supplementary Figure S1)**. As the fish developed, the beads stayed in the remaining yolk, close to the yolk extension, still permitting magnetic orientation of the larva. The injected embryos could be rotated in a non-contact manner within their chorion by moving a permanent magnet past the sample (**Supplementary video S1)**. This rotation arises from the attraction of the injected beads by the permanent magnet. The applied force led to a slight translation and a rotation of the embryo, minimizing the distance between the beads and the magnet.

To have a dynamic control of the magnetic field we used electromagnets for the following experiments. As the force applied on a superparamagnetic bead is proportional to the magnetic field gradient ^8^, we sharpened the core of the electromagnets to create a sufficiently strong magnetic field gradient even with a moderate current (300 mA), keeping the heating of electromagnets to a minimum. To orient the sample in three dimensions, we used four magnets arranged in a tetrahedral geometry around the sample in the center **(Figure 1b)**. An injected zebrafish embryo was embedded in an FEP tube. The inner diameter (1.0 mm) was slightly smaller than the chorion (∼1.2 mm) to prevent the fish from being pulled out of the tube by the lowest magnet **(Figure 1c)**. By sequentially switching between four magnets, we could rotate the embryo in a non-contact manner and position it in four different orientations given by the tetrahedral arrangement of electromagnets **(Figure 1c)**.

To study any sample in its optimal orientation, intermediate positions can be accessed in two different ways: spherical samples with no preferred orientation (zebrafish embryo until the tail-bud stage) can be stopped during rotation from one magnet to the next by switching the current off. Alternatively, by applying currents to two or more magnets simultaneously, the embryo orients along the resulting magnetic field. Hence, by changing the ratio of the currents applied to the two magnets, the embryo can be rotated continuously from one magnet to the other and positioned at any orientation between the two magnets **(Figure 1d)**.

We found that the rotation of the zebrafish embryo from one magnet to another (corresponding to 109.5°) took less than 30s at a current of 300 mA **(Figure 1e)**. This transition time was inversely proportional to the current and could therefore be tuned according to the application **(Supplementary Figure S2)**. After releasing the embryo from the magnetic field, it retracted in less than 10 s before settling into its final position. The embryo remained stable for over a minute, sufficient to acquire a 3D stack of the whole embryo on a SPIM system **(Figure 1e)**.

To image developing zebrafish in its optimal orientation with high resolution and low photo-toxicity, we implemented the tetrahedral electromagnet configuration around the sample chamber in a SPIM setup **(Figure 2a).** Typically, SPIM provides only a single axis of rotation for multi-view imaging ^3,9^. We asked if the additional degrees of freedom in our setup could lead to improved coverage of the sample.

**Figure 2.**
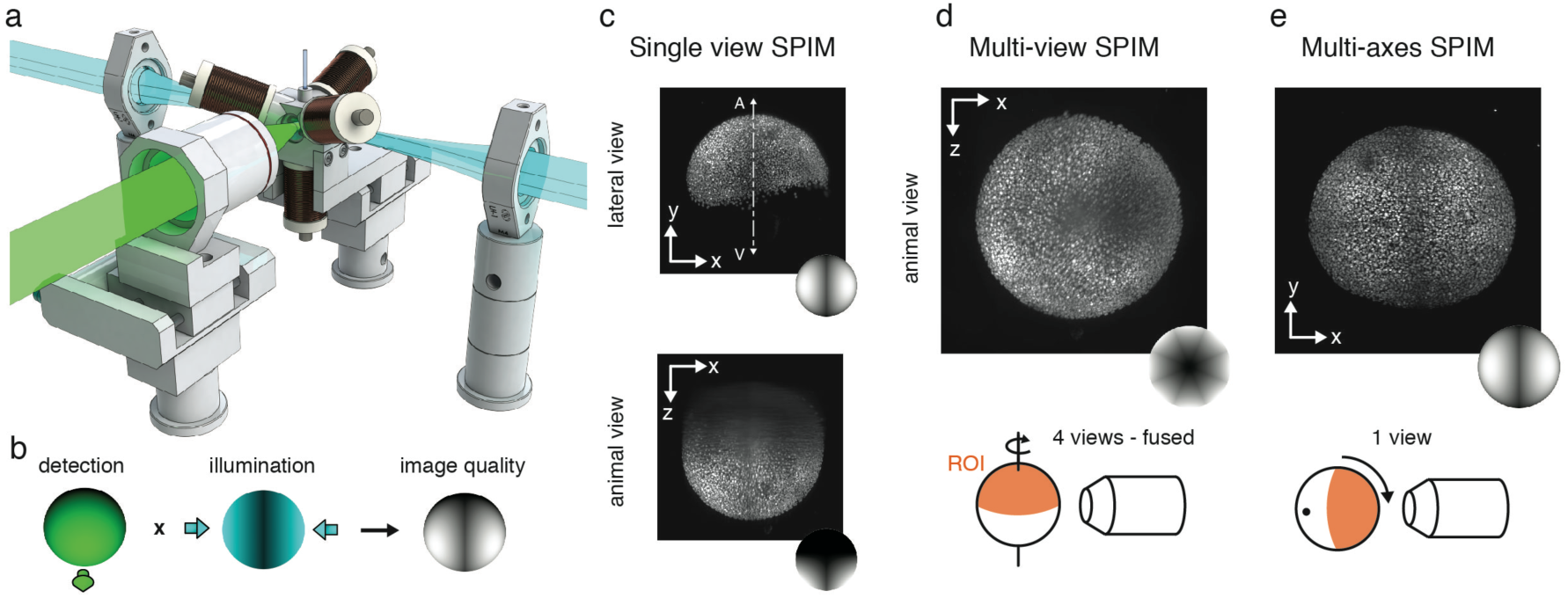
Non-contact 3D orientation technique adapted to a SPIM system. (***a***) Schematic of the SPIM setup with the tetrahedral electromagnets arranged around the sample tube. The magnets were held by the sample chamber and the sample was illuminated by a light sheet through two windows. The fluorescence was detected through a third window by a detection objective. The sample chamber and the detection objective were motorized to move the sample through the light sheet and to correct for the different path lengths in air and water. (***b***) Schematic representation of the obtained image quality in SPIM imaging, which is influenced by the sample orientation relative to the detection and illumination objectives. (***c***) Schematic representation of the image quality obtained by a single view on a conventional SPIM system. Only the lateral view shows high optical quality, whereas the image quality in the view on the animal pole shows the decreasing image quality along z. (***d***) Schematic representation of the image quality obtained by multi-view SPIM imaging for four views (0°, 45°, 180° and 225°). The fusion of the four views shows a homogenous resolution along the equator but a decreasing resolution towards the cap (rotational axis). (***d***) Schematic representation of the image quality obtained with the multi-axes SPIM technique. The view on the animal pole shows the isotropic image quality of the animal pole (ROI) obtained by orienting the animal pole towards the detection objective by the multi-axes sample orientation technique.

In multi-view imaging, the view directly facing the detection objective is well detected but the image quality suffers from oblique illumination. At the same time, the orthogonal views facing the illumination objectives are well illuminated but poorly resolved due to longer optical path. Therefore, the best image quality is achieved in the region between the illumination and detection objectives, which is well detected and well illuminated ^10^ **(Figure 2b/c)**. When rotating the sample in a conventional, single rotational-axis system, uniform resolution is achieved only along the equator. However, the image quality at the poles is always poor owing to its inaccessibility for illumination and detection.

We imaged an injected zebrafish embryo (*Tg(h2afva:h2afva-GFP)* ^11^) in the conventional multi-view SPIM mode and in the new multi-axes SPIM mode (**Figure 2d/e**). During early development, the animal pole is on top of the yolk (north pole) and therefore only poorly accessible for the conventional SPIM since the rotational axis of the SPIM is parallel to the animal-vegetal axis (A-V-axis) (**Figure 2c**). To still reconstruct the animal pole, we imaged the sample in four different orientations by rotating the sample tube, registering and fusing these non-ideal views. (**Figure 2d)**.

In contrast, in the multi-axes SPIM mode we were able to orient the animal pole facing the detection objective and a single stack was sufficient to image the entire region of interest (ROI) (**Figure 2d**). The resolution was superior and no longer limited by the inferior axial resolution as in the conventional reconstruction. Thus, by positioning the sample in its ideal orientation we achieve superior resolution compared to multi-view SPIM with less views needed.

Traditional single-lens light microscopes do not offer any sample rotation and can image the sample from only one side. Our magnetic sample rotation is instrumental in this case as well. We developed an insert consisting of a plate and an arc holding two electromagnets **(Figure 3a)**, which can be easily adapted to any commercial light microscope. As a proof of concept, we show two-view imaging of a zebrafish larva (5dpf, *Tg(kdrl:GFP)*^12^) on an upright epi-fluorescence microscope. The magnet orientation can be adapted continuously by sliding the magnets along the arc. We found that the rotation of the zebrafish larva worked best with an angle of about 35° between the electromagnet and the plate.

**Figure 3.**
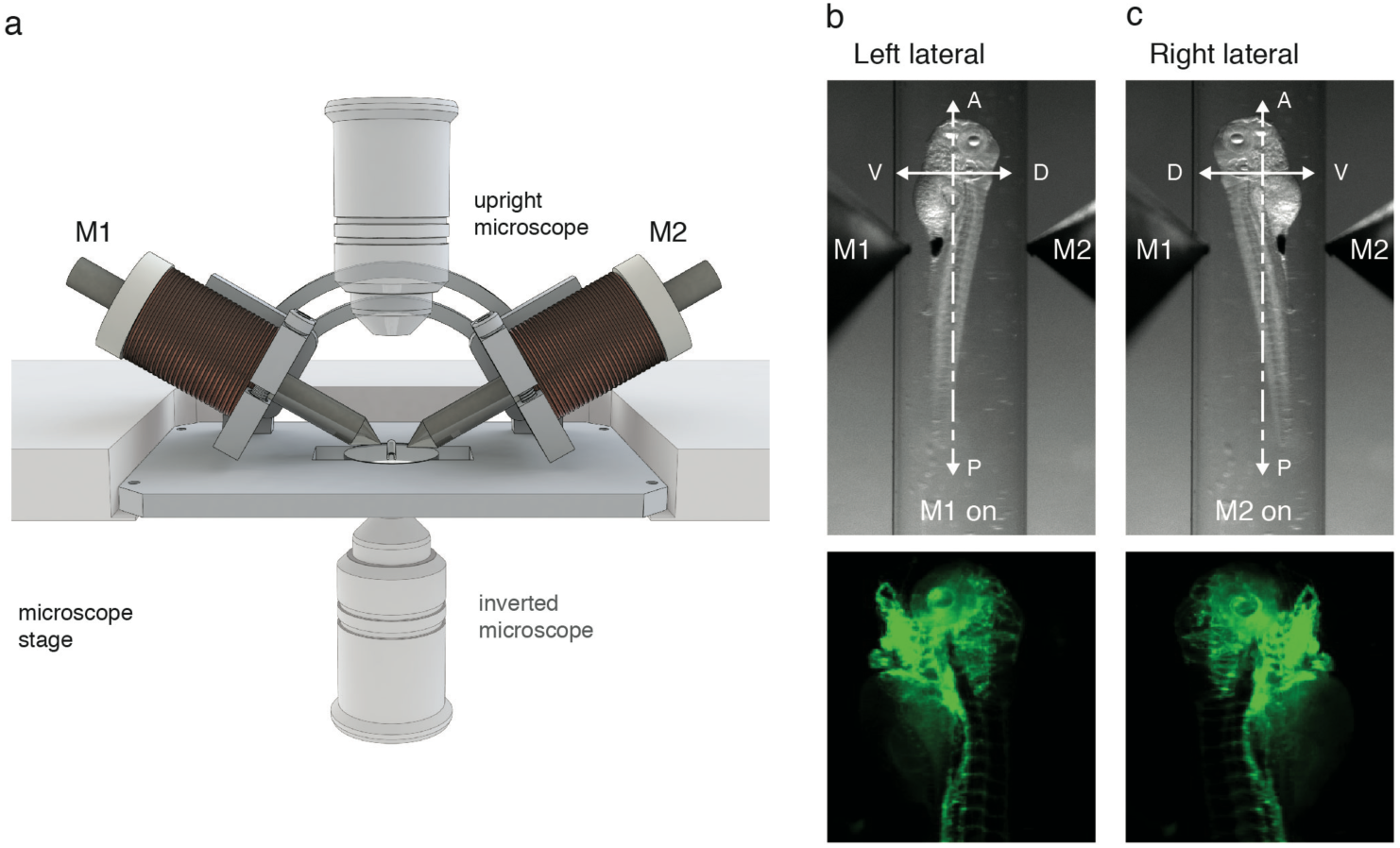
Non-contact 3D orientation technique adapted to a commercial epi-fluorescence microscope. (***a***) Schematic showing the inset holding two electromagnets (M1 and M2) on a microscope stage. (***b,c***) Bright-field and fluorescence images of a 5dpf Tg(kdrl:GFP) zebrafish larva rotated about its anterior-posterior axis by powering electromagnet M1 and M2, respectively.

The zebrafish larva was embedded in a glass capillary and positioned in the intersection of the magnet axes. We oriented the larva about its anterior-posterior axis. By switching on one magnet, the larva was slightly translated towards the wall of the glass capillary and rotated about 180° towards the magnet. By switching the magnet off, the larva was released from the force and settled in its resting position **(Figure 3 b/c)**. A 180-degree rotation from the left lateral to the right lateral resting position took no longer than 10 s with 1 A current, illustrating the that our system can add multi-view capabilities to any conventional microscope.

We have developed a non-contact method to orient specimens in a microscope by injecting them with superparamagnetic beads and applying an external magnetic field. We optimized the protocol for zebrafish embryos and larvae and adapted the method to a custom-built SPIM and a commercial epi-fluorescence microscope to enable multi-axes, multi-view imaging.

Our technique relies on magnetic forces applied on beads injected into an organism and therefore the following points need to be considered. First, high magnetic forces can lead to the deformation of the sample or a translation of the beads within the yolk, which could potentially damage the sample. We prevented these effects by using only minimal forces for a short time. Second, the magnetic forces can lead to a slight translation of the sample within the chorion for zebrafish embryos and within the glass capillary for zebrafish larva, respectively. However, translation is anyway necessary to center the region of interest after rotation.

We optimized our technique for zebrafish embryos and larvae. For the embryo, we made use of the fact, that the embryo is encapsulated in its fluid filled chorion facilitating the rotation; for the larva, we exploited its elongated shape. Other model organisms with a less suited morphology could benefit from our technique by embedding them in a sphere or cylinder of agarose that sits in a fluid filled volume.

High-throughput methods^13^, which currently do not offer any rotational axis for sample orientation, will also greatly benefit from our non-contact sample orientation technique. All samples can be put in the same orientation, increasing the screening sensitivity and specificity. Photo-manipulation techniques benefit especially from the full flexibility to orient the sample such that the intervention can be precisely done in the desired region.

## Materials and Methods

### SPIM setup

The SPIM setup consisted of an Olympus N4X-PF air objective for detection and two lenses (AC254-060-A, Thorlabs, Germany) aligned orthogonal to the detection for double sided light-sheet illumination. The sample was illuminated alternately from two sides. The alternating illumination was controlled via a flipper mirror (8892-K-M, New Focus, USA). The sample chamber was 3D-printed and had two windows for illumination and one window for detection of the sample. The electromagnets were held by the sample chamber and oriented in a tetrahedral geometry. The applied current was remotely controlled via a programmable power supply (QL355P, Aim-TTi, United Kingdom) and a manual switch. The focus of the four electromagnet-tips coincided with the sample. The sample was inserted from the top within an FEP tube orthogonal to the detection objective and the illumination lenses. For sample scanning the sample chamber was mounted on a motorized linear stage (M111.1DG, Physik Instrumente, Germany) and moved relative to the detection objective. To correct for the different path length in water and air the detection objective was also placed on a linear stage (M111.1DG, Physik Instrumente, Germany) and moved accordingly. For wide-field illumination a LED (CCS TH-27/27-SW, Stemmer imaging GmbH, Germany) was placed behind the sample. For light-sheet illumination a single color (488nm) Coherent Sapphire laser (488-30 CDRH) was used and the light-sheets were generated with cylindrical lenses (ACY254-050-A, Thorlabs, Germany). The wide-field and fluorescence signals were detected with a sCMOS camera (Zyla 5.5, Andor, United Kingdom).

### Electromagnets

The electromagnets were custom-built and consisted of a magnetic core made from HyMu 80 alloy (National Electronic Alloys, USA) and a solenoid (**Supplementary Figure S3).** The bobbin was made from Teflon and a high resistance wire was wound up hundreds of times to create a high magnetic field. The magnetic core had a diameter of 6 mm and was tapered to create a magnetic field gradient. The inner diameter of the bobbin was 6 mm in which the magnetic core could be inserted.

### Superparamagnetic beads

The washed superparamagnetic beads (Dynabeads MyOne Carboxylic Acid, Invitrogen, USA) with a diameter of 2.8 µm were injected with standard glass needles into the yolk of the zebrafish embryo. The injection needle was inserted from either the vegetal pole or the lateral side. For a sufficient torque to turn the embryo it is important that the beads stay close to the yolk membrane. Injections were performed at very low pressure (5-10 psi) and long injection duration (100-150 ms) to avoid the dispersion of beads. Since the beads sink to the tip of the needle and change the concentration of the bead solution the pressure had to be increased for some injections (∼45psi) to prevent blocking of the needle. By applying a strong constant magnetic field with a permanent magnet, after the injection of the beads, the beads are attracted and clumped. This aggregation of beads prevents the single beads from translating through the yolk and eases the rotation.

### Magnetic manipulator inset for a stereoscope and inverted microscopes

The magnetic manipulator insert was built from an aluminum plate with a size of 92.5x64 mm fitting on standard microscope stages. On this plate, an arc was mounted, holding two electromagnets that can be freely slid along the arc to control the angle between the magnets and the sample. The sample was either embedded in a glass capillary for mounting on an upright microscope or placed on a coverslip and held by a Teflon holder to study the sample on an inverted microscope.

### Zebrafish

Zebrafish (*Danio rerio*) adults and embryos were kept at 28.5 °C and were handled according to established protocols ^14^. Transgenic lines *Tg(kdrl:GFP)* ^12^ and *Tg(h2afva:h2afva-GFP)* ^11^ were used. For SPIM imaging, the zebrafish embryos were either embedded in small (inner diameter 1.0 mm, outer diameter 1.6 mm) FEP tubes in E3 or in bigger (inner diameter 1.6 mm, outer diameter 2.4 mm) FEP tubes in 1.5% low-melting-point agarose (Sigma).

For imaging the zebrafish larvae on the stereoscope, the embryos were treated with 0.2mM 1-phenyl 2-thiourea (Sigma) at 24h post fertilization to inhibit melanogenesis. During imaging, the samples were anaesthetized with 200 mg/l Tricaine (Sigma) and embedded inside glass capillaries in E3 containing 200 mg/l Tricaine. All animals were treated in accordance with EU directive 2011/63/EU as well as the German Animal Welfare Act.

### Sample handling for taking multi-view and multi-axes stacks

We embedded the zebrafish embryo (*Tg(h2afva:h2afva-GFP)*) in a FEP tube (inner diameter 1.6 mm, outer diameter 2.4 mm) in 1.5% low-melting-point agarose (Sigma). For taking the multi-view SPIM data we manually rotated the sample tube. We took two views 45° apart from both sides of the sample ^10^. The four different angles (0°, 45°, 180° and 225°) were registered and fused with the feature-based registration using the nuclei as features ^15,16^.

For taking the multi-axes stack we oriented the same injected embryo within its chorion towards the detection objective with the four electromagnets. Before recording a stack, we switched the magnet off to avoid any deformations by the applied force. Switching off resulted in a slight reorientation in a new settle position.

After a few seconds, when the embryo had settled in its new resting position, a 3D stack of the embryo was acquired.

### Kymograph of rotating zebrafish embryo

To characterize the rotation of the zebrafish embryos, we generated kymographs of the bright field videos showing their reorientation **(Figure 1e and Supplementary Figure S2)**. Since the embryos undergo rotation we generated kymographs by slicing in time along the path of the magnetic beads using FIJI ^17^. This path was drawn manually and kept for comparing different currents **(Supplementary Figure S2).**

## Funding

Work in the lab of J.H. was supported by the Max Planck Society and European Research Council (CoG SmartMic, 647885)

## Competing Financial interests

G.S. and J.H. are inventors on a patent application (PCT/EP2016/057186), which is related to the described research.

## Author Contributions

G.S. and J.H. conceived the idea. F.B. developed instrumentation; F.B., G.S. and J.H. designed experiment; F.B. and G.S. performed experiments and analyzed data; F.B. and J.H. wrote the manuscript, all authors discussed results and critically revised the manuscript.

## Supplementary Figures

**Figure S1.**
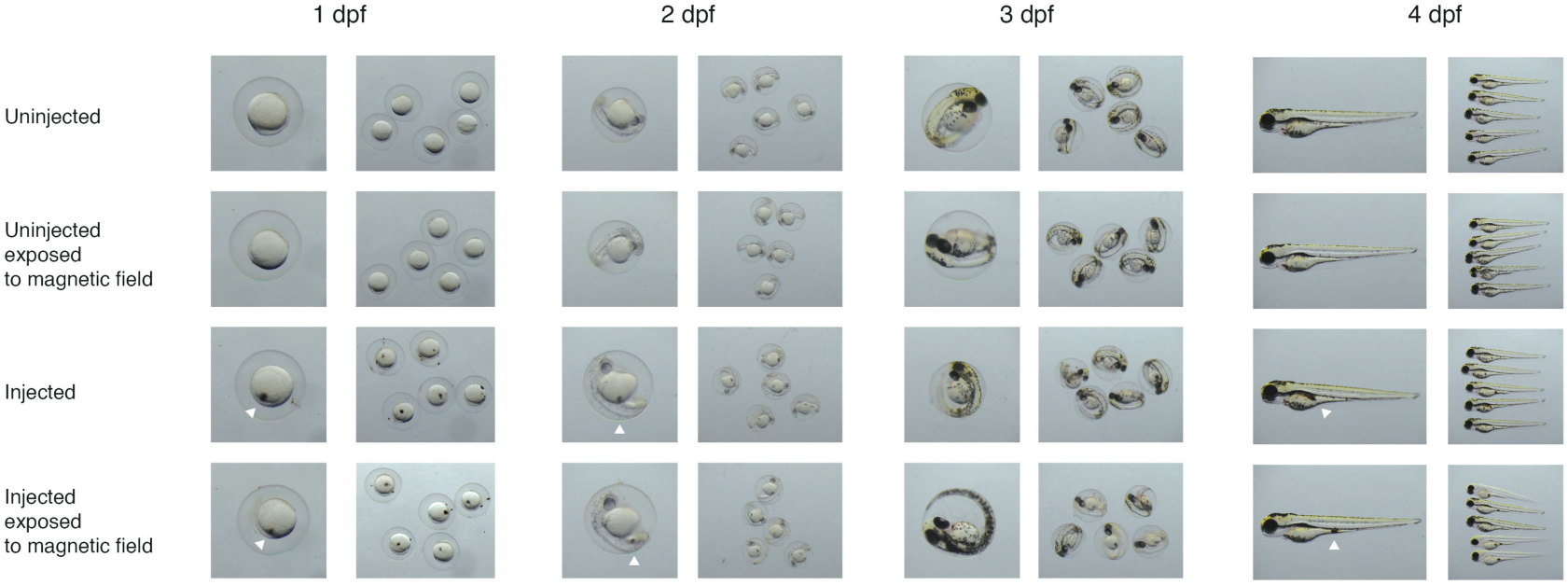
Bright-field images of injected and uninjected zebrafish embryos and larvae over four days. The embryos have been injected according to the ***Supplementary protocol 1*** with 2.8 µm beads (white arrow heads). Injected and uninjected zebrafish have been exposed to magnetic field by placing a permanent magnet close to the dish. All controls showed no delay in development.

**Figure S2.**
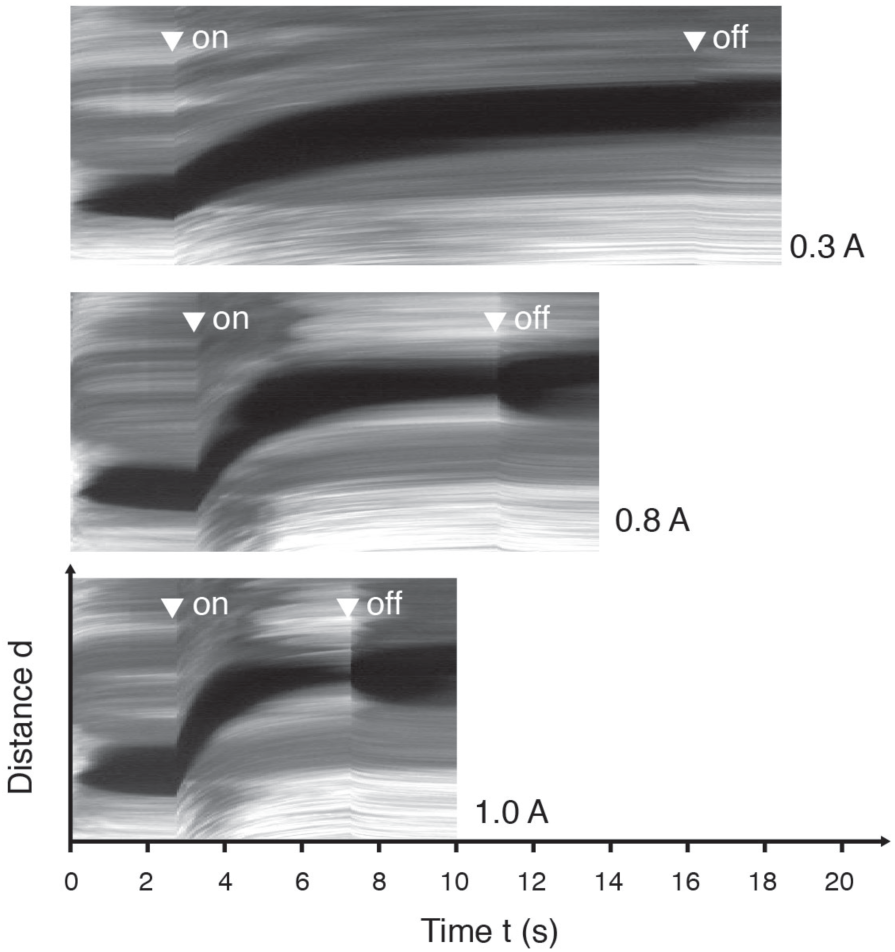
Rotation of an injected zebrafish with different currents showing that the rotational speed can be tuned by the applied current. Arrow heads indicate when the electromagnet was switched on and off, respectively.

**Figure S3.**
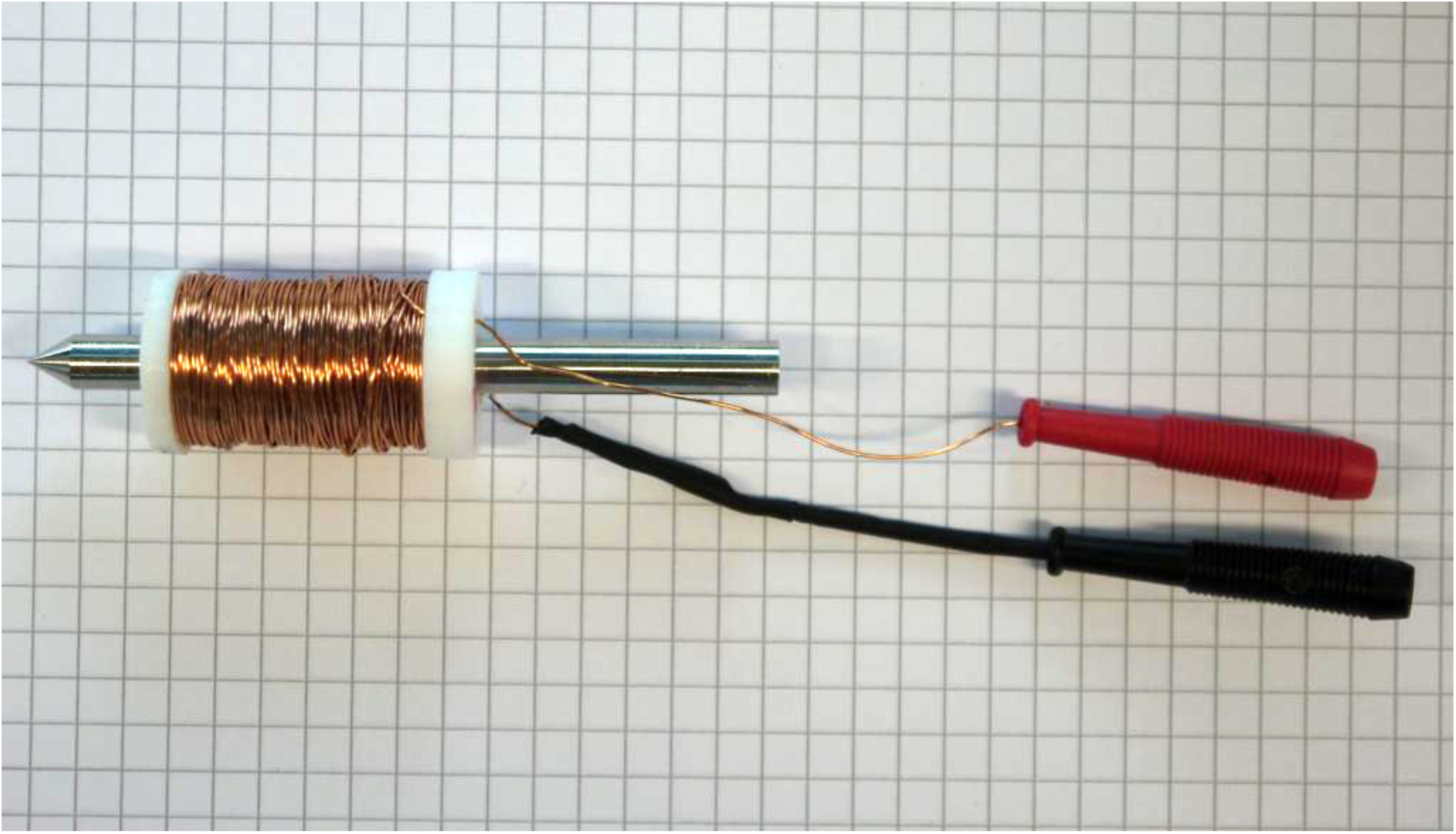
Photograph of one of the four custom-built electromagnets. Shown is the sharpened core inside a Teflon bobbin with the copper coil. Two cables connect to the power supply.

## Supplementary Movies

**Movie S1**: 4hpf zebrafish embryo oriented by permanent magnet.

**Movie S2:** 5hpf embedded in FEP tube and oriented by electromagnets in SPIM setup.

**Movie S3:** 5dpf zebrafish larva in glass capillary oriented by electromagnets on epi-fluorescence microscope.

## Supplementary Information

### Protocol 1: Washing and injection of superparamagnetic beads

#### Step 1: Washing the beads

Take 10μl of the stock bead solution (10mg/ml) in an Eppendorf tube. Bring a permanent magnet close to the tube to clump the beads and remove the remaining solution using a pipette. Take the magnet away and re-suspend the beads in 20μl of distilled water. Repeat the process twice before injecting the diluted bead solution (5mg/ml) into embryo.

#### Step 2: Microinjection

Injections were performed using a micro-injector and injection needles. The opening of the needle is adjusted such that it is not too small for the beads to come out as well as not too large to damage the embryo. Beads are injected at extremely low pressure (∼10psi) and long injection duration (∼150ms) in order to avoid dispersion of beads in the yolk. Since the beads sink to the tip of the needle and change the concentration of the bead solution, the pressure had to be increased for some injections (∼45psi) to prevent blocking of the needle.

#### Step 3: Aggregation of injected beads

After injecting the beads, a strong constant magnetic field is applied with a permanent magnet to attract and clump the beads. This aggregation of beads preserves the single beads from translating through the yolk and eases the rotation.

#### Step 4a: Embedding zebrafish embryo for SPIM experiments

The zebrafish embryo in E3 buffer is sucked up into a FEP tube with a syringe. To prevent the embryo from being pulled out of the tube the embryo is either embedded in E3 in a tube with an inner tube diameter (1.0 mm), which is a bit smaller than the zebrafish chorion (1.2 mm) or in a bigger tube (inner diameter 1.6 mm) in 1.5% low-melting-point agarose (Sigma). In both cases the embryo (0.8 mm) is still free to move within its chorion.

#### Step 4b: Embedding zebrafish larvae (5 dpf) for epi-fluorescence microscope experiments

The fish larva with 200 mg/l Tricaine (Sigma) in E3 is sucked into a glass capillary and can be exposed to the magnetic field to rotate it about its anterior-posterior axis.

